# Epidermal cells sculpt sensory nerve endings through actomyosin contractility

**DOI:** 10.64898/2026.08.28.747813

**Authors:** Min Lee, Julie Underwood, Jing Xu, Ru-Rong Ji, Terry Lechler

## Abstract

Peripheral sensory neurons innervate the skin to detect mechanical, thermal, and noxious stimuli. Within the epidermis, nerve fibers terminate beneath tight junctions, shielding them from environmental exposure. Although epidermal differentiation coordinates tight junction assembly, its role in organizing nerve terminals is poorly understood. Here, we show that activation of Notch, a master regulator of epidermal differentiation, caused near-complete loss of epidermal innervation. This was largely the result of increased contractility rather than impaired differentiation. Inducing epidermal contractility was sufficient to deplete nerve fibers with striking spatial precision, and restoring normal contractility reversed this effect. Actomyosin contractility is highest in the granular layers of the epidermis, where tight junctions form and nerve fibers terminate. Ablation of nonmuscle myosin II allowed nerve fibers to extend beyond their normal termination zone and caused touch hypersensitivity. Together, these findings demonstrate that epidermal contractility positions sensory nerve endings through spatially controlled pruning and defines a mechanical boundary established by epidermal cells that restrict neuronal outgrowth.

## Introduction

The skin serves not only as a physical barrier against environmental insults but also as a sensory organ that enables the detection of thermal, mechanical, and chemical stimuli. The ability of the skin to rapidly sense the environment is essential for organisms to avoid danger and acts as a driver of social development (Nascimento et al., 2018). This sensory system also underlies cutaneous pain and itch, major clinical pathologies.

In mice and human, cutaneous sensory information is transmitted by dorsal root ganglion (DRG) neurons, whose peripheral axons detect stimuli in the skin and whose central axons convey these signals to the spinal cord (Meltzer et al., 2021; Nascimento et al., 2018). Distinct populations of DRG neurons innervate the epidermis where they intimately interact with resident keratinocytes (Erbacher et al., 2024; Qi et al., 2024; Talagas et al., 2020a; Talagas et al., 2020b). Separated from the dermis by a layer of basement membrane, the epidermis is composed of a basal progenitor layer, which proliferates and gives rise to differentiated suprabasal cells, called spinous cells (Moreci and Lechler, 2020; Prado-Mantilla and Lechler, 2023). These, in turn, differentiate into granular cells, which form tight junctions (TJs) that demarcate the upper boundary of epidermal innervation (Takahashi et al., 2019). TJ function within the granular cells depends on high levels of actomyosin contractility that is unique to these epidermal cells (Rübsam et al., 2017; Sumigray et al., 2012). Notably, keratinocytes are constantly turning over in the epidermis, necessitating neuronal endings to remodel and form new associations with these cells.

Keratinocytes are integral components of the sensory system that play essential roles in directing innervation and mediating sensation (Talagas et al., 2020a; Yang and Chien, 2019; Yin et al., 2021). These epidermal cells modulate neuronal morphology through several mechanisms. For example, keratinocyte-secreted nerve growth factor (NGF) promotes axonal growth and epidermal targeting (Albers et al., 1994; Davis et al., 1997). In addition, specialized keratinocytes (Krt17 positive) are required for innervation and function of sensory structures called touch domes (Jenkins et al., 2019). Further, distinct DRG neurons target different differentiated layers within the epidermis. Nonpeptidergic MrgprD (Mas-related G protein-coupled receptor D) neurons innervate the granular layer, whereas peptidergic CGRP (calcitonin gene-related peptide) neurons target the spinous layer (Wang and Zylka, 2009; Zylka et al., 2005). This layer-specific termination of nerve endings suggests a keratinocyte-dependent control of epidermal nerve patterning. Finally, in adult mouse and human skin, sensory nerve endings are precisely maintained beneath tight junctions in the granular layers (Takahashi et al., 2019). In patients with atopic dermatitis, this local pruning does not occur and may contribute to the development of chronic itch (De Benedetto et al., 2011; Takahashi et al., 2019; Yuki et al., 2016). These studies suggest that homeostatic pruning in the granular layers is crucial for maintaining nerve endings beneath tight junctions and regulating sensory signaling. However, the mechanisms regulating this local pruning remain unclear. Together, these data highlight the myriad ways that keratinocytes communicate with and regulate neuronal morphology and function. Where understood, these effects are mediated through chemical signals, either secreted factors or through direct cell-cell contacts.

In this study, we find that the mechanical status of keratinocytes has a profound effect on neuronal innervation. We initially set out to understand how epidermal differentiation regulates the precise targeting of distinct neuronal subtypes in the epidermis. We found that preventing granular cell differentiation by activating Notch signaling unexpectedly led to an almost total loss of epidermal innervation. RNA-seq analysis revealed that a contractile gene signature was upregulated in Notch-activated epidermis. Using both pharmacologic rescue and direct genetic perturbation of contractility, we found that epidermal contractility locally and rapidly induces neuronal severing. Reducing epidermal contractility by ablating non-muscle myosin proteins caused nerve fibers to overextend into the most superficial epidermis and resulted in touch hypersensitivity. This study demonstrates that the mechanical properties of epidermal cells are a key regulator for maintaining and organizing sensory nerve endings in the epidermis.

## Results

### Activated Notch signaling in keratinocytes disrupts epidermal innervation

During development, DRG sensory neurons extend their peripheral axons as bundled nerve fibers, migrating from the DRG into the skin (Maklad et al., 2009; Nascimento et al., 2018; Peters et al., 2002). To examine epidermal innervation in developing and adult murine skin, we visualized the nerve fibers by staining DRG neurons with a pan-neuronal marker PGP9.5 (Protein Gene Product 9.5, also known as *UCH-L1*). As previously reported (Peters et al., 2002), sensory axons reach the basement membrane at embryonic day 17.5 (E17.5), where they are found exclusively in the dermis in both back hairy skin and paw non-hairy skin (hereafter referred as glabrous skin) (Supplementary Fig.1A-D). By E18.5, DRG neurons begin to cross the basement membrane and progressively innervate the epidermis. Notably, epidermal innervation occurs at the same developmental stage across anatomically distinct skin types, although how neurons breach the basement membrane and coordinate innervation across the body are not known. At postnatal day 0 (P0), axons continue to extend and branch, eventually forming a mature arborization by adulthood. In adult mice, nerve fibers follow a more circuitous path within the thinner epidermis of back skin. In contrast, neuronal axons project more linearly through the thicker epidermis and terminate in the upper granular layer in adult glabrous skin.

A functional skin barrier relies on the formation of tight junctions in granular layers and the establishment of the cornified envelope, which develop shortly before sensory neurons begin to innervate the epidermis (Furuse et al., 2002; Hardman et al., 1998; Sevilla et al., 2013). We therefore tested whether this terminal differentiation was required for neurons to enter the epidermis. We used conditional activation of Notch signaling in the epidermis (Krt14-CRE;ROSA-NICD; this allows the active Notch intracellular domain, NICD, to be expressed in basal progenitor cells), which is reported to drive spinous cell fate and results in loss of granular layer markers (Blanpain et al., 2006). The GFP is expressed bicistronicly with NICD and serves as a marker for recombination (Fig. 1B). Consistently, we found that constitutively activating Notch signaling resulted in decreased levels of the granular marker, loricrin (Supplementary Fig. 2A and 2B). We examined innervation in E18.5 back skin by staining for PGP9.5. In wild-type embryos, nerve fibers were observed in both the dermis and the epidermis (Fig. 1A and 1C). Surprisingly, in NICD-expressing epidermis, innervation was severely disrupted, with an almost complete loss of IENFs (intra-epidermal nerve fibers). Despite the loss of epidermal innervation, nerve fibers remained in the dermis, demonstrating that Notch activation specifically impairs epidermal targeting rather than overall axonal pathfinding in the skin.

**Figure 1.**
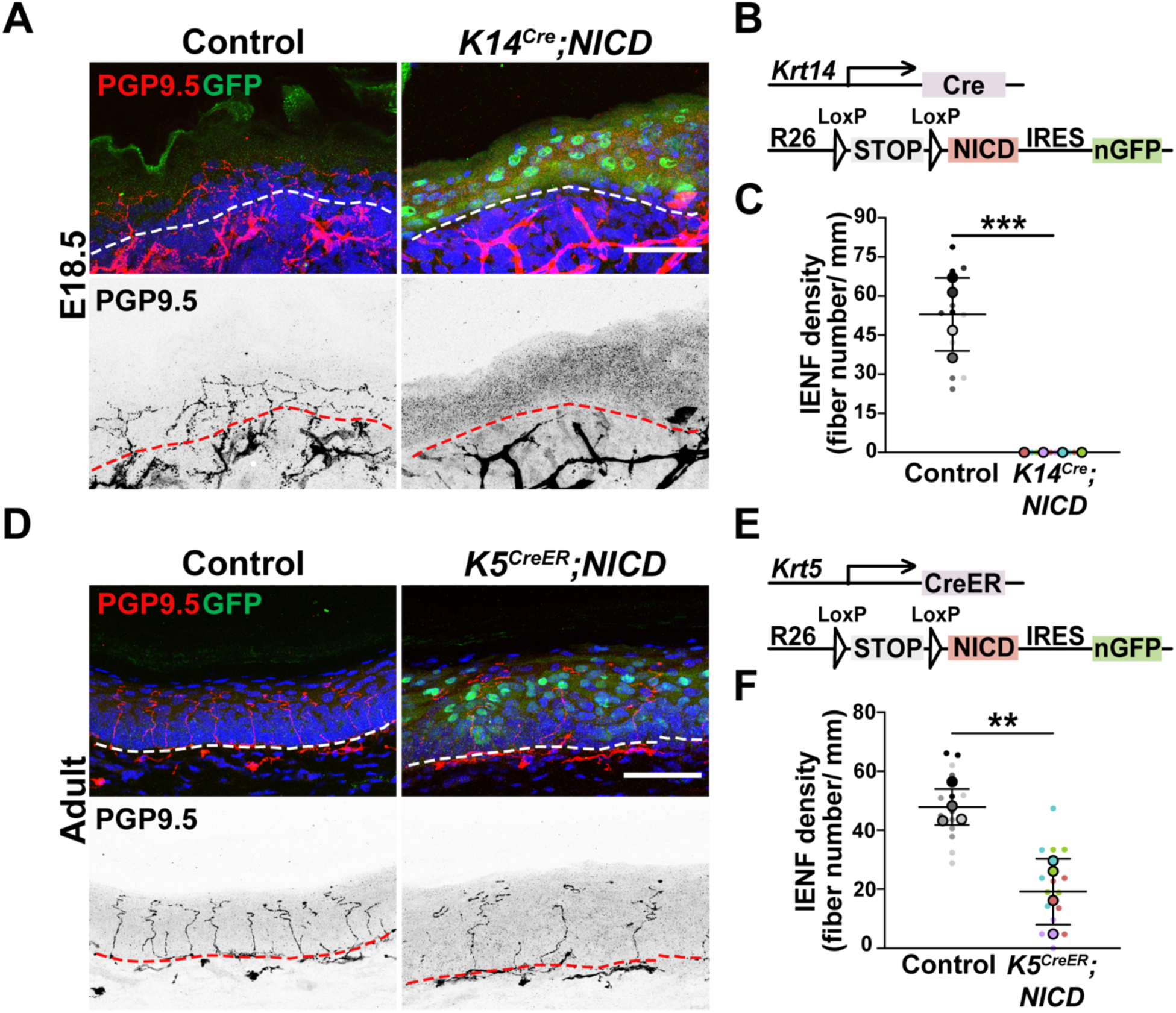
Activated Notch signaling in keratinocytes disrupts epidermal innervation. (A) Immunofluorescence staining of neurons (PGP9.5 in red; inverted image below), NICD-expressing keratinocytes (GFP in green) and nuclei (blue) in control and K14-NICD back skin at E18.5. Dashed lines: basement membrane. (B) Diagram of the K14^Cre^;NICD mouse model. (C) Quantification of IENF density in E18.5 control and K14-NICD back skin. n = 4 embryos per group, 3-4 fields per embryo. Data are mean ± SD. p = 0.003, unpaired t-test. (D) Immunofluorescence staining of neurons (PGP9.5 in red; inverted image below), NICD-expressing keratinocytes (GFP in green) and nuclei (blue) in control and K5^CreER^;NICD adult paw non-hairy skin. Mice were intraperitoneally administered with tamoxifen for 2 days and collected 5 days after the final injection. (E) Diagram of the K5^CreER^;NICD mouse model. (F) Quantification of IENF density in control and K5^CreER^;NICD adult epidermis in paw non-hairy skin. n = 4 animals per group, 4 fields per animal. Data are mean ± SD. p = 0.0041, unpaired t-test. Scale bars: 50 μm.

In this mouse model, Notch is activated before epidermal innervation occurs. Therefore, it was unclear whether neurons failed to innervate or whether they were disrupted after entering the epidermis. To address this, we induced NICD expression in the basal progenitor cells of adult mice (Krt5-CRE^ER^;ROSA-NICD)(Fig. 1E), in which epidermal innervation was already established. We administered tamoxifen intraperitoneally for two consecutive days to induce recombination and examined innervation 5 days after the final injection. The thin epidermis in adult back skin makes it difficult to visualize and quantify nerve fibers. Thus, we assessed epidermal innervation in glabrous skin. NICD expression in adult epidermis resulted in a significantly reduced IENF density compared to control mice (Fig. 1D and 1F). In contrast to the embryonic mice, the fluorescence intensity level of loricrin was comparable between control and Krt5-CRE^ER^;ROSA-NICD mice (Supplementary Fig. 2C and 2D), indicating that granular markers are not lost in this time frame. These findings demonstrate that inducing Notch signaling in keratinocytes disrupts epidermal innervation in both embryonic and adult skin, and suggests that signaling downstream of NICD, but not loss of terminal differentiation, contributes to this phenotype.

### Notch activation increases epidermal contractility

To investigate the mechanisms underlying the innervation defects in NICD-expressing epidermis, we isolated epidermal cells from control and K14-CRE;ROSA-NICD back skin at E17.5 for bulk RNA-seq analysis. Principal component analysis (PCA) demonstrated that samples segregated by their genotype (Supplementary Fig. 3A). Gene Ontology (GO) analysis of the 1,791 genes down-regulated in NICD-expressing epidermis revealed enrichment of downregulated biological processes related to epidermal differentiation, tight junction assembly, and barrier formation (Blanpain et al., 2006; Sumigray et al., 2014) (Fig. 2A-B). This is consistent with previous studies showing that Notch activation promotes spinous cell fate while blocking granular differentiation, resulting in differentiation defects and hence early postnatal death (Blanpain et al., 2006). Analysis of the upregulated genes, in contrast, revealed an unexpected increase of transcripts encoding proteins involved in actin cytoskeleton regulationand actomyosin organization (Fig. 2C).

**Figure 2.**
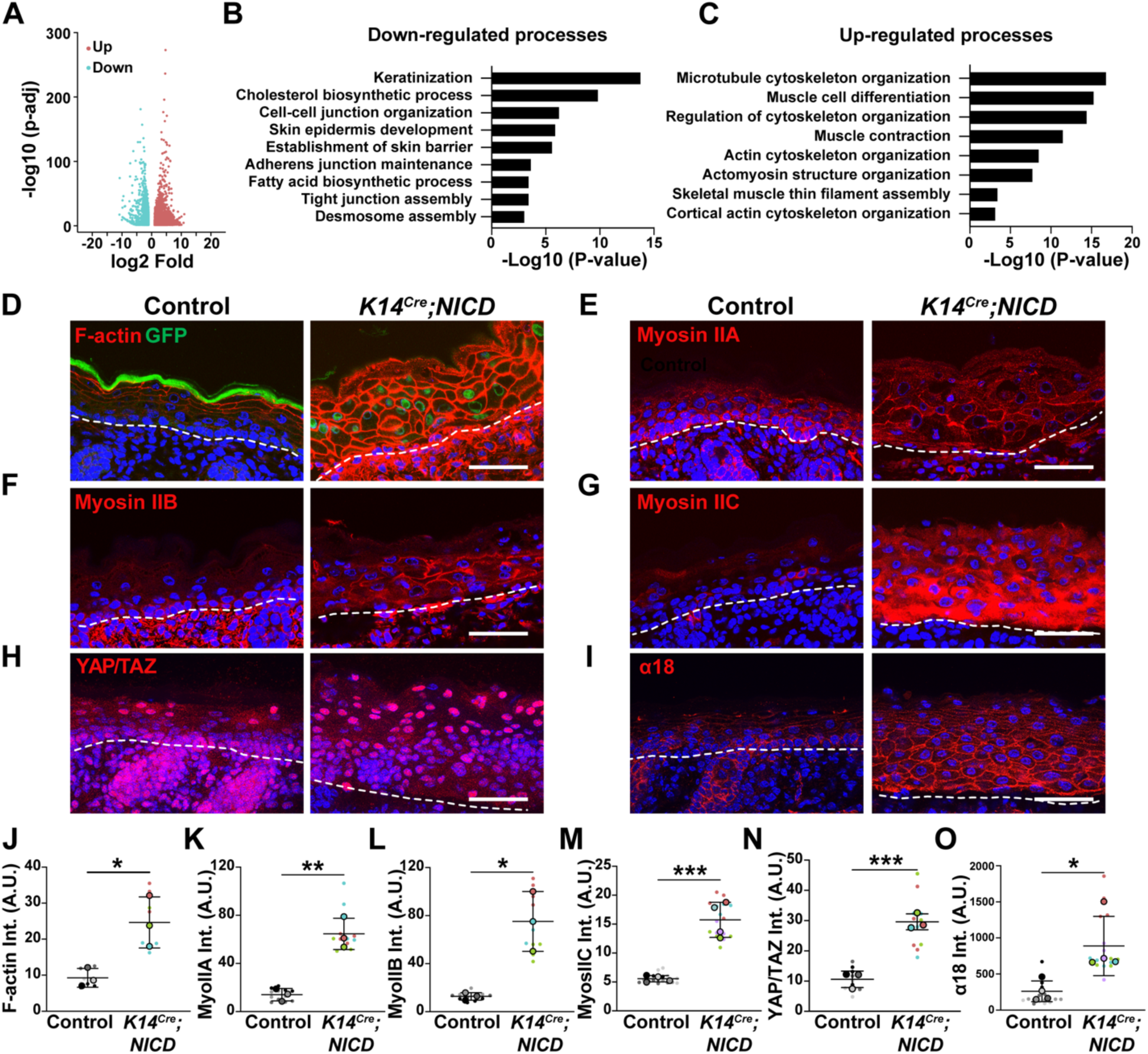
Contractility signature genes are up-regulated in NICD-expressing epidermis. (A) Volcano plot of differentially expressed genes from RNA-seq data comparing K14-NICD (Notch activated) and WT epidermal cells. n=3 per genotype. (B and C) Selected top biological process GO terms for up-and down-regulated genes identified in E17.5 control and NICD-expressing epidermis. (D) Co-staining of GFP (green) and F-actin (red) in control and NICD-expressing E18.5 back skin. Dashed lines: basement membrane. (E-G) Immunofluorescence staining of non-muscle myosin proteins (myosin IIA, IIB, and IIC) in control and K14-NICD mice at E18.5. Dashed lines: basement membrane. (H) Immunofluorescence staining of YAP/TAZ (red) and (I) α-18 (red) in E18.5 control and K14-NICD back skin. (J) Quantification of F-actin fluorescence intensity in control and K14-NICD epidermis. n = 3 embryos per group, 3 fields per embryo. Data are mean ± SD. p = 0.0243, unpaired t-test. (K) Quantification of cortical myosin IIA fluorescence intensity of epidermal cells. n = 3 embryos per group, 4 fields per embryo. Data are mean ± SD. p = 0.0033, unpaired t-test. (L) Quantification of cortical myosin IIB fluorescence intensity of epidermal cells. n = 3 embryos per group, 4 fields per embryo. Data are mean ± SD. p = 0.0126, unpaired t-test. (M) Quantification of myosin IIC fluorescence intensity in control and K14-NICD epidermis. n = 4 embryos per group, 4 fields per embryo. Data are mean ± SD. p = 0.0006, unpaired t-test. (N) Percentage of suprabasal cells with nuclear YAP/TAZ in E18.5 control and K14-NICD back skin. n = 4 embryos per group, 3-4 fields per embryo. Data are mean ± SD. p = 0.001, unpaired t-test. (O) Quantification of cortical α-18 fluorescence intensity of epidermal cells. n = 4 embryos per group, 4 fields per embryo. Data are mean ± SD. p = 0.0279, unpaired t-test. Scale bars: 50 μm.

To better understand how these transcriptional changes manifest in tissue, we examined cell morphology and epidermal architecture by staining F-actin with fluorescent phalloidin to visualize the cell cortex. Mutant epidermis exhibited a dramatic increase in cortical F-actin across epidermal layers as compared with control animals (Fig. 2D and 2J). The increased levels of F-actin extended even into the dermis, although the underlying cause of this non-cell autonomous response remains unknown. We also found that epidermal cells became more rounded with a reduced cell aspect ratio (Fig. 2D and Supplementary Fig. 3B-C), a morphological change associated with the increased cortical F-actin and actomyosin contractility (Ning et al., 2021). Not only was cortical F-actin increased, but we also found increased levels of three type II myosins, myosin IIA, IIB, and IIC (Fig. 2E-G and Fig. 2K-M). The mechanoresponsive transcriptional co-activator YAP/TAZ (Dupont et al., 2011) provided another readout of cellular contractility in epidermal cells, with prominent nuclear localization in contractile cells (Ning et al., 2021; Prado-Mantilla et al., 2025). The percentage of suprabasal cells exhibiting nuclear YAP/TAZ was significantly increased in NICD-expressing epidermis compared with controls (Fig. 2H and 2N). Furthermore, we stained for α18, a marker that recognizes a tension-sensitive epitope of the adherens junction protein α-catenin (Yonemura et al., 2010). Quantification showed increased levels of cortical α18 in K14-Cre;ROSA-NICD mice, suggesting that junctional tension is elevated upon Notch activation in the epidermis (Fig. 2I and 2O). Together, these results demonstrate that Notch activation increases cellular contractility and junctional tension in the epidermis, which may contribute to the disruption of epidermal innervation.

### Loss of canonical Notch signaling in keratinocytes does not impair epidermal innervation in embryonic skin

Epidermal-specific conditional deletion of RBP-J, a transcriptional mediator of Notch signaling, inhibits canonical Notch signaling and impairs epidermal differentiation during development (Blanpain et al., 2006). Ablation of RBP-J in the epidermis did not significantly alter innervation, as IENF density was comparable between K14-CRE;RBP-J^f/f^ and control embryos (Supplementary Fig. 3D and 3E). However, consistent with previous studies (Blanpain et al., 2006), these mice had a thinner epidermis with reduced spinous layers (Supplementary Fig. 3F-G). While loricrin intensity remained unchanged in the granular layers (Supplementary Fig. 3G), mice died from dehydration after birth. There were no changes in cortical F-actin intensity in RBP-J deficient epidermis (Supplementary Fig. 3I-J). These results suggest that canonical Notch signaling is not necessary for establishing epidermal innervation in embryonic skin, although Notch overactivation is sufficient to disrupt innervation.

### Increased epidermal contractility disrupts IENFs in embryonic and adult skin

To directly test whether increasing epidermal contractility causes nerve fiber loss, we took advantage of a mouse line, TRE-Arhgef11^CA^, that allows genetically inducible actomyosin contractility (Ning et al., 2021). The protein expressed is a constitutively active form of the Rho-guanine nucleotide exchange factor Arhgef11, tagged with HA. Its expression increases cellular contractility through activation of the Rho/ROCK/myosin II pathway (Berlew et al., 2021; Valon et al., 2017). We crossed TRE-Arhgef11^CA^ to K10-rtTA mice (hereafter referred to as K10-Arhgef11) to induce actomyosin contractility in differentiated epidermis (Fig. 3A). K10-Arhgef11 embryos exhibited a thickened epidermis, and an almost complete loss of epidermal innervation in the back skin (Fig. 3B and 3D). We were able to take advantage of the occasional mosaic expression of K10-Arhgef11 to assess how locally it acted. In areas devoid of Arhgef11 expression, sensory neurons crossed the basement membrane and innervated the epidermis (Fig. 3C and 4B, arrowheads). This finding indicates that nerve fibers are selectively excluded from contractile regions, and this is a very local effect. Increased contractility also disrupted epidermal innervation when induced in adult back and paw skin (Fig. 3C, 3E and 4B-E), demonstrating that increased contractility impairs epidermal innervation across developmental stages and skin types. To determine whether this phenotype resulted from elevated epidermal contractility, we treated adult mice with Y-27632 to inhibit ROCK signaling upstream of actomyosin contractility (Uehata et al., 1997). Epidermal innervation was partially rescued by Y-27632 treatment, resulting in IENF density similar to control animals (Fig. 3F-H). Further, in NICD-expressing epidermis, Y-27632 treatment also partially rescued epidermal innervation at E18.5 (Fig. 3I-L). Together, these findings demonstrate that epidermal contractility is a key regulator of nerve innervation.

**Figure 3.**
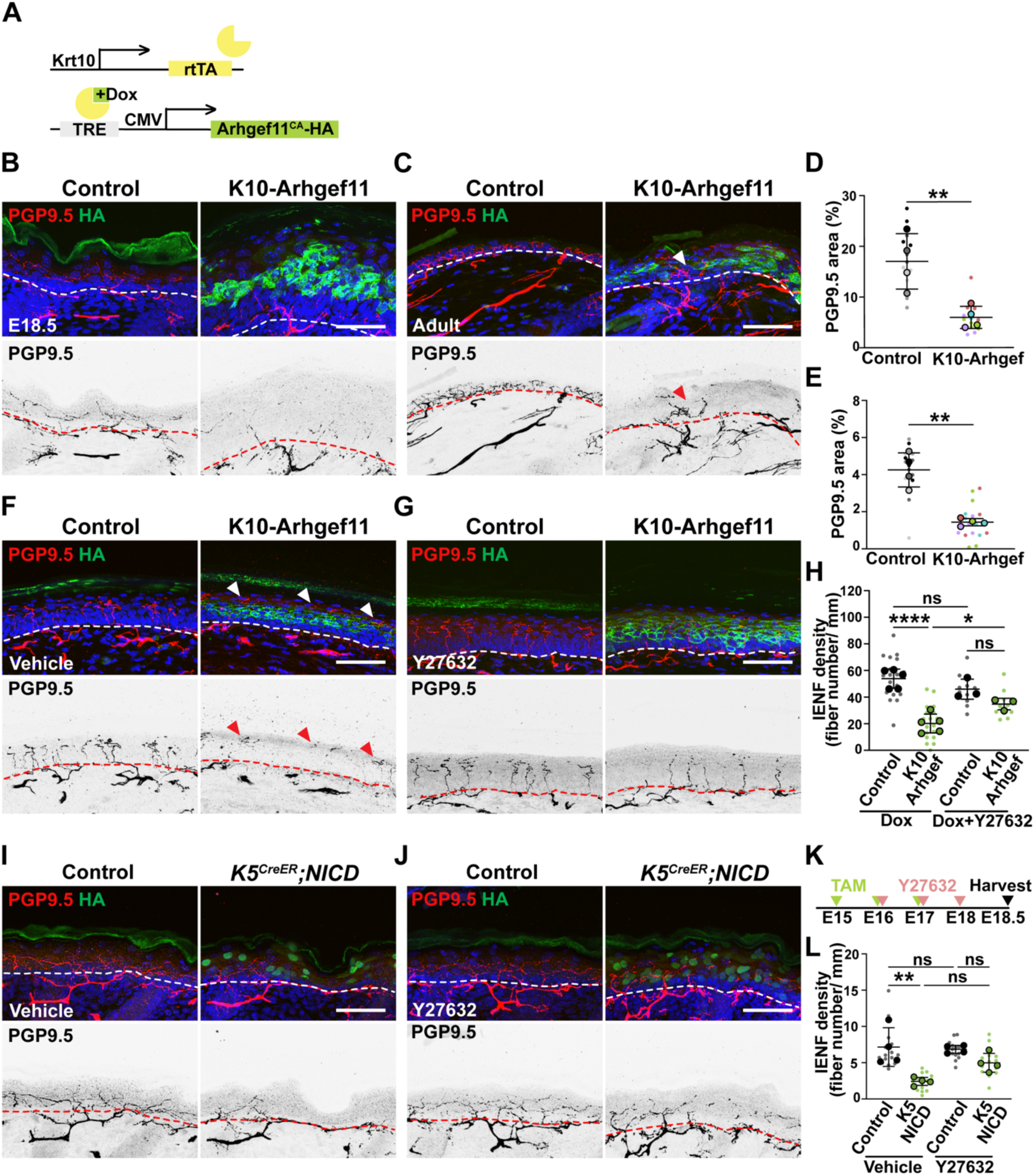
Increased epidermal contractility disrupts innervation in embryonic and adult skin. (A) Diagram of the K10-rtTA;TRE-Arhgef11^CA^ mouse model. (B-C) Immunofluorescence staining of neurons (PGP9.5 in red; inverted image below), Arhgef11^CA^-HA (green) and nuclei (blue) in control and K10-Arhgef11 back skin at E18.5 (B) and in adults (C). Green signal in the cornified envelope at the top of epidermis is autofluorescence. (D-E) The percentage of PGP9.5^+^ nerve area in the epidermis of back skin at E18.5 (D, n = 4 embryos per group, 4 fields per embryo. Data are mean ± SD. p = 0.0053, paired t-test), and in adult (E, n = 4 mice per group, 3-4 fields per animal. Data are mean ± SD. p = 0.0093, unpaired t-test). (F-G) Immunofluorescence staining of neurons (PGP9.5, red), Arhgef11^CA^-HA (green) in control and K10-Arhgef11 adult paw glabrous skin. Mice were treated with doxycycline together with vehicle or Y27632 for 12 hours. (H) Quantification of IENF density in F and G. n = 3-5 mice per group, 3-5 fields per animal. Data are mean ± SD. ns = not significant, *p = 0.0499, ****p < 0.0001, one-way ANOVA analysis. (I,J) Immunofluorescence staining of nerve fibers (PGP9.5, red), nucleus GFP (green) in control and K5^CreER^;NICD back skin at E18.5 (Y27632-treated and untreated mice). (K) Diagram of the Y27632 rescue experiment in K5^CreER^;NICD mice. (L) Quantification of the percentage of PGP9.5^+^ nerve area in E18.5 epidermis. n = 4 embryos per group, 3-4 fields per embryo. Data are mean ± SD. ns = not significant, **p = 0.0037, one-way ANOVA analysis. Scale bar: 50 μm.

**Figure 4.**
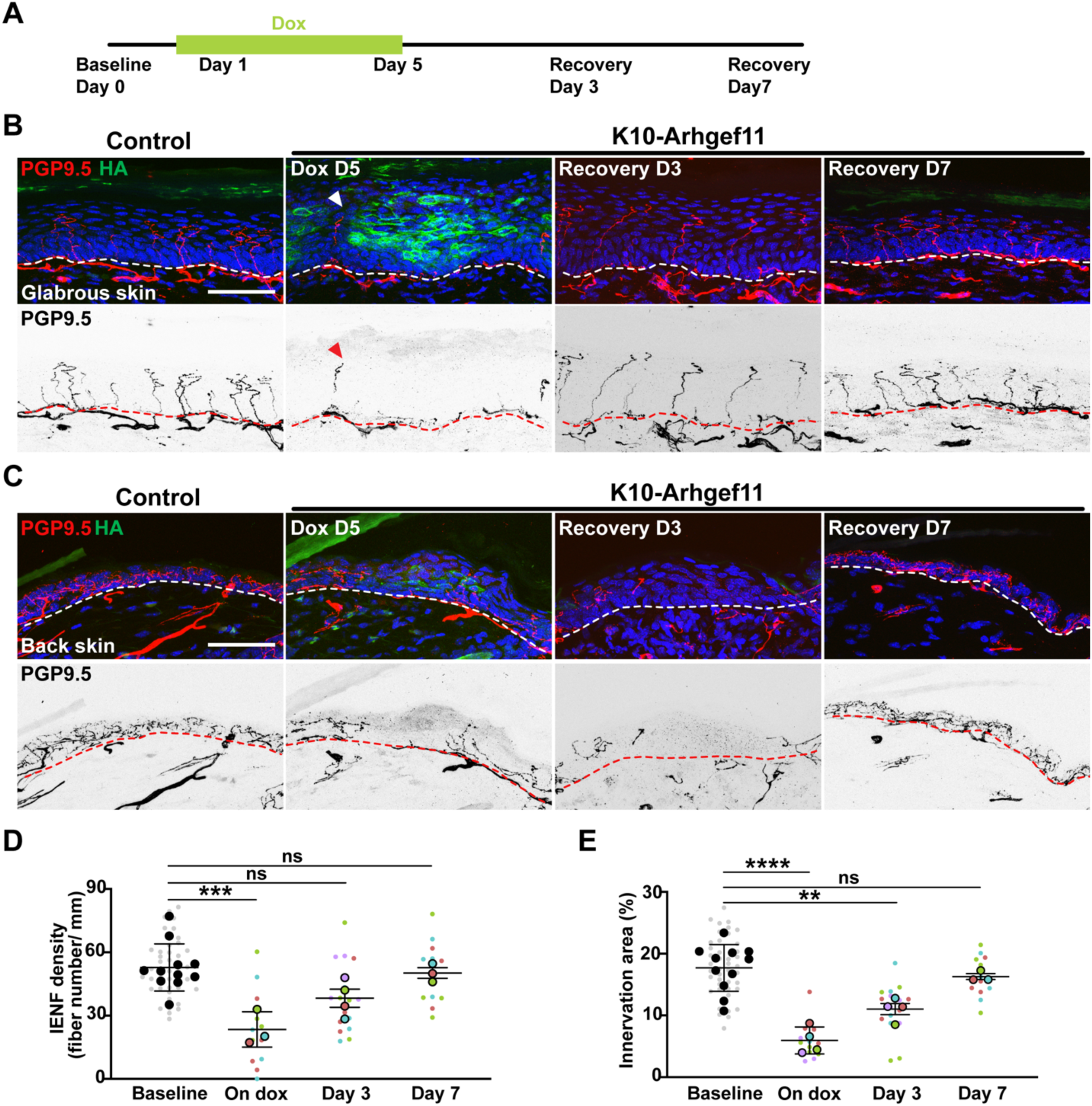
Sensory neurons reinnervate the epidermis upon restoration of normal contractility. (A) Diagram of the doxycycline treatment regimen for examining epidermal nerve fibers during severing and regeneration. Mice were fed dox food from P50-P55 to induce epidermal contractility and switched back to normal mouse chow after P55 for the recovery phase. (B,C) Immunofluorescence staining of neurons (PGP9.5, red), and Arhgef11^CA^-HA (green) in control and K10-Arhgef11 adult glabrous skin (B) and back skin (C). Dashed lines: basement membrane. Scale bar: 50 μm. (D) Quantification of IENF density in paw non-hairy skin of control and K10-Arhgef11 mice. n = 3-4 mice per group, 3-4 fields per animal. Data are mean ± SD. ns = not significant, *p = 0.0359, ***p = 0.0008, one-way ANOVA. (E) The percentage of PGP9.5^+^ nerve area in the epidermis of control and K10-Arhgef11 adult back skin. n = 3-4 animals per group, 3-4 fields per animal. Data are mean ± SD. ns = not significant, **p = 0.0046, ****p < 0.0001, one-way ANOVA.

### Restoration of epidermal innervation following normalization of contractility

Unlike most of the neurons in the central nervous system, peripheral sensory neurons have a remarkable capacity to regenerate following injuries (Mahar and Cavalli, 2018; Varadarajan et al., 2022). However, functional recovery is often limited due to axonal misguidance or aberrant sprouting of neighboring uninjured nerves (Gangadharan et al., 2022; Jeon et al., 2024). In the spared nerve injury (SNI) model, nociceptive neurons reinnervate the denervated skin but fail to invade the epidermis. Lack of intraepidermal nerve free endings is a pathological hallmark of several painful neurological disorders, including diabetic peripheral neuropathy (Hu et al., 2023; Tian et al., 2024) and chemotherapy-induced peripheral neuropathy (Wozniak et al., 2018). Our K10-Arhgef11 mice provide a model for specifically injuring epidermal nerve fibers and examining their roles in sensory function. To address whether sensory neurons regenerate their epidermal nerve fibers after the contractility-induced depletion, we replaced dox chow with regular mouse food to restore contractility to normal physiological levels and allowed the epidermis to recover (Fig. 4A). Within 3 days, epidermal nerve fibers had re-innervated the epidermis with normal levels of innervation within a week (Fig. 4B-E). The regeneration kinetics differed slightly between back skin and glabrous skin with back skin showing a lower reinnervation capacity at 3 days post recovery.

Because our data showed that epidermal nerve fibers were severed in contractile epidermis within 12 hours after doxycycline induction (Fig. 3F-H), we first asked whether the rapid depletion of epidermal nerve terminals was sufficient to cause sensory dysfunction. Mechanical sensitivity, measured by applying von Frey filaments to hind paw skin, was not altered despite the loss of epidermal nerve fibers (Supplementary Fig. 4A-B) at this stage (day 1). We observed no difference in thermal withdrawal latency and cold response duration between K10-Arhgef11 and control animals (Supplementary Fig. 4C-D), indicating that thermal sensitivity also remains unchanged. These results suggest that acute IENF loss is not sufficient to perturb mechanical-or thermal-evoked pain responses, though the mechanisms that might underlie this robustness are not known. Following 5 days of increased epidermal contractility (day 5), mechanosensitivity was decreased, though it is unclear whether this is a primary effect of loss of epidermal innervation or is secondary to increased epidermal thickness. In contrast, mice exhibited elevated thermal sensitivity to both heat and cold stimuli, which may result from an inflammatory response with increased infiltration of immune cells in the dermis. During epidermal reinnervation, mechanical hypersensitivity improved at day 3 and recovered to baseline after a week.

### Epidermal contractility is required for nerve pruning at tight junctions and normal touch sensation

Nociceptive nerve fibers are pruned and retained beneath tight junctions in granular layers, protecting them from direct exposure to the environment (Bäsler et al., 2016; Takahashi et al., 2019). Importantly, our previous work demonstrated that epidermal contractility is required for tight junction barrier function (Sumigray et al., 2012), and additional studies demonstrated that contractility is highest in the granular layers (Miroshnikova et al., 2018; Rübsam et al., 2017). We therefore asked whether genetically decreasing epidermal contractility affects nerve fiber pruning in the granular layers. We first generated K5-CRE^ER^;Myh9^fl/fl^ mice (hereafter referred to as MyoIIA cKO) to reduce contractility in keratinocytes by genetically ablating the gene that encodes non-muscle myosin IIA. Myosin IIA is the major type II myosin in the epidermis by RNA-Seq analysis (Joost et al., 2016). In adult glabrous skin, myosin IIA was detected throughout the epidermis in control mice, and its levels were significantly reduced after 5 days of depletion in MyoIIA cKO mice (Supplementary Fig. 5A, 5B, 5G and 5I). Ablation of myosin IIA resulted in notable changes in epidermal nerve fibers (Fig. 5A and 5C). There was a small decrease in epidermal innervation, and the nerve fibers that did innervate showed altered paths, being less linear and more parallel to the basement membrane. In addition, the nerve fibers in MyoIIA cKO mice terminated more superficially in the epidermis compared with controls, as evidenced by a shorter distance between the tip of nerve terminals and the outermost epidermal layer (Fig. 5D). Moreover, the thickness of loricrin^+^ epidermis was not changed compared with control animals, suggesting that the superficial termination of nerve fibers in Myosin IIA cKO epidermis is unlikely due to granular differentiation defects (Fig. 5E). To further reduce contractility, we generated K5-CRE^ER^;Myh9^fl/fl^;Myh10^fl/fl^ mice (hereafter referred to as Myosin IIA/B dKO) to ablate both non-muscle myosin IIA and IIB in the epidermis. Immunohistochemistry and western blot analysis revealed that both myosin IIA and IIB levels were significantly reduced (Supplementary Fig. 5C-5F, 5G, and 5I). Myosin IIA/B dKO mice exhibited lower IENF density, less linear nerve fibers, aberrant superficial termination (Fig. 5F and 5H-I), and intact granular layers in the epidermis (Fig. 5G and 5J). The similarity to the MyoIIA cKO mice suggests that myosin IIA is a major contributor to the contractility in the epidermis.

**Figure 5.**
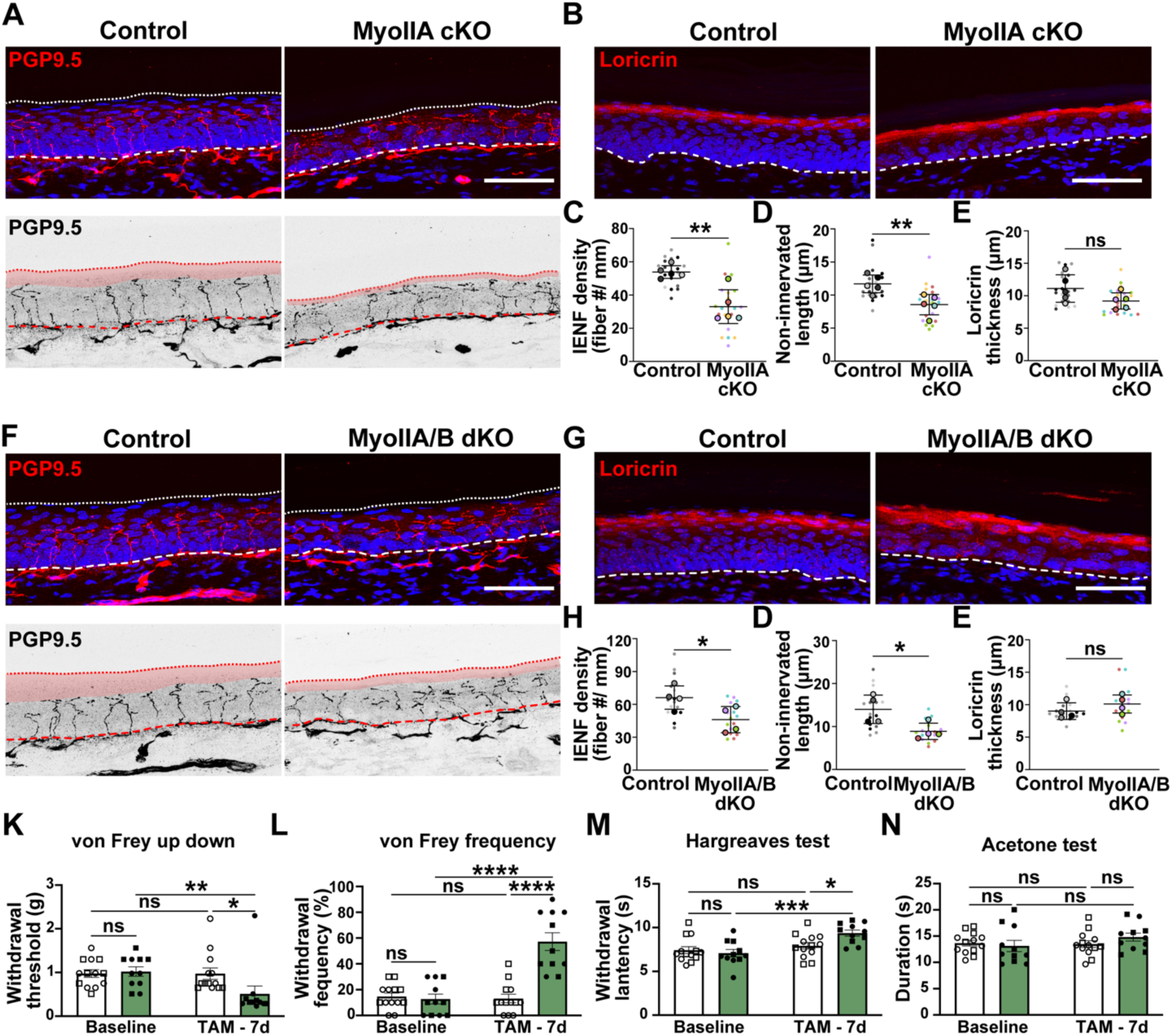
Reduced epidermal contractility leads to superficial nerve termination and mechanical hypersensitivity. (A) Immunofluorescence staining of neurons (PGP9.5, red) and nuclei (blue) in control and K5^CreER^;Myh9 adult paw glabrous skin. Dashed lines: basement membrane. Dotted lines: the most superficial epidermal layer. The area between nerve terminals and the outermost epidermal layer is highlighted in red in the inverted images. (B) Immunofluorescence staining of loricrin (red) in control and K5^CreER^;Myh9 adult paw glabrous skin. (C) Quantification of IENF density in control and K5^CreER^;Myh9 paw non-hairy skin. n = 4 mice per group, 4 fields per animal. Data are mean ± SD. p = 0.0457, unpaired t-test. (D) Quantification of the distance between nerve terminals and the top epidermal layer in (A). n = 4 mice per group, 4 fields per animal. Data are mean ± SD. p = 0.0358, unpaired t-test. (E) Quantification of the thickness of loricrin+ epidermal layers in control and K5^CreER^;Myh9 paw non-hairy skin. n = 4 mice per group, 4 fields per animal. Data are mean ± SD. ns = not significant, unpaired t-test. (F) Immunofluorescence staining of neurons (PGP9.5, red) in control and myosin 2A/B dKO adult paw non-hairy skin. (G) Immunofluorescence staining of loricrin (red) in control and myosin 2A/B dKO adult paw glabrous skin. (H) Quantification of IENF density in control and myosin 2A/B dKO paw non-hairy skin. n = 4 mice per group, 4 fields per animal. Data are mean ± SD. p = 0.0457, unpaired t-test. Scale bars: 50 μm. (I) Quantification of the distance between nerve terminals and the top epidermal layer in (F). n = 4 mice per group, 4 fields per animal. Data are mean ± SD. p = 0.0358, unpaired t-test. (J) Quantification of the thickness of loricrin+ epidermal layers in control and myosin 2A/B dKO paw non-hairy skin. n = 4 mice per group, 4 fields per animal. Data are mean ± SD. ns = not significant, unpaired t-test. (K and L) von Frey tests in Myosin 2A/B dKO mice. n =11-13 per group. Data are mean ± SEM. *p = 0.0149, **p = 0.0094, ***p = 0.0002, ****p < 0.0001, ns = not significant, two-way ANOVA followed by Sidak post hoc test. Circle, male; square, female. (M) Hargreaves tests in myosin 2A/B knockout mice. Data are mean ± SEM. *p = 0.0112, ***p = 0.0003, ns = not significant, two-way ANOVA followed by Sidak post hoc test. Circle, male; square, female. (N) Acetone tests in myosin 2A/B knockout mice. n =11-13 per group. Data are mean ± SEM. ns = not significant, two-way ANOVA followed by Sidak post hoc test. Circle, male; square, female.

Defective pruning of epidermal nerve fibers exposes their terminals to the environment and hence sensitizes sensory neurons (Takahashi et al., 2019). To determine whether the aberrant superficial termination of nerve fibers affects cutaneous sensations, we performed behavioral tests to evaluate mechanical and thermal sensitivity. Compared with control animals, Myosin IIA/B dKO mice exhibited mechanical hypersensitivity, a slight reduction in heat sensitivity, and no change in cold responses (Fig. 5K-N). Consistent with these findings, touch sensation is the major modality mediated by MrgprD neurons, whose axons extend up to the granular layer (Cavanaugh et al., 2009; Qi et al., 2024; Wang and Zylka, 2009; Zylka et al., 2005). The lack of substantial temperature sensitivity further indicates this is not simply a generalized sensitization of cutaneous neurons to stimuli. These findings demonstrate that epidermal contractility is required for proper nerve pruning in the granular layers and normal skin touch sensitivity.

### A local increase in epidermal contractility severs nerve fibers

Our data suggest that myosin-dependent contractility in the granular cells forms a mechanical barrier that prevents nerve extension. To further test this, we established the K14-rtTA;TRE-Arhgef11^CA^ mouse line (hereafter called K14-Arhgef11 mice) to induce contractility in basal keratinocytes, effectively moving this barrier lower in the epidermis (Fig. 6A,E). Strikingly, as early as 6 hours post-induction, epidermal nerve fibers were severed within the basal layer where contractility was increased (Fig. 6B). Quantification revealed a significantly reduced number of nerve fibers crossing the first epidermal layer (Fig. 6EC. Although the number of nerve fibers in the upper non-contractile epidermis was slightly reduced, most nerve fibers remained detectable (Fig. 6D). By 24 hours post-induction, the remaining nerve fiber fragments in the suprabasal epidermis were lost (Supplementary Fig. 6A-C). Contractility could work directly on neurons or could induce local formation of tight junctions that result in nerve severing. We therefore asked whether elevated contractility induces the ectopic formation of tight junctions. Staining for the tight junction protein ZO-1 showed no ectopic expression in basal cells (Supplementary Fig. 6D), indicating that the local nerve pruning resulted from increased cellular contractility rather than tight junction formation. Together, these data demonstrate that epidermal cells spatially regulate sensory nerve endings by contractility-dependent pruning (Fig. 6E).

**Figure 6.**
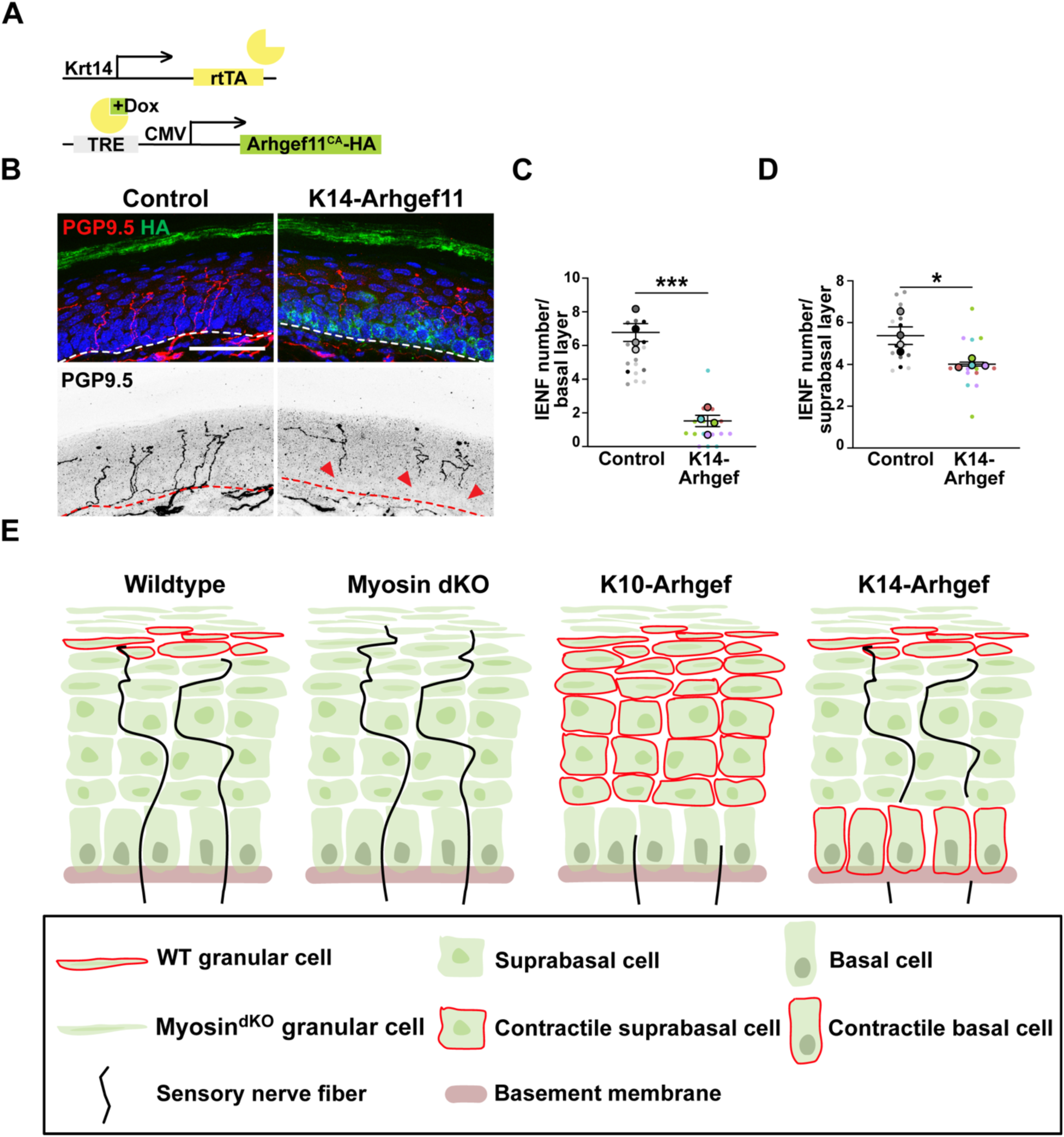
Increasing contractility in epidermal cells results in rapid and local nerve pruning. (A) Diagram of the K14-rtTA;TRE-Arhgef11^CA^ mouse model. (B) Immunofluorescence staining of neurons (PGP9.5 in red), contractile basal keratinocytes (HA in green) and nuclei (blue) in control and K14-Arhgef11 adult glabrous skin. Doxycycline treatment: 6 hours. Dashed lines: basement membrane. Arrowheads: nerve severance. Scale bar: 50 μm. (C) Number of nerve fibers crossing basal layer in control and K14-Arhgef11 adult glabrous skin, at 6-hour post doxycycline treatment. n = 4 mice per group, 4 fields per animal. Data are mean ± SD. p = 0.0002, unpaired t-test. (D) Number of nerve fibers remained within suprabasal layers in control and K14-Arhgef11 adult glabrous skin, at 6-hour post doxycycline treatment. n = 4 mice per group, 4 fields per animal. Data are mean ± SD. p = 0.0199, unpaired t-test. (E) Summary of results, indicating nerve fiber position relative to areas of epidermal contractility.

## Discussion

This work identifies a non-cell-autonomous mechanism by which epidermal contractility controls sensory neuron morphology. Keratinocytes are known to provide chemical signals that promote sensory axon growth into the epidermis (Albers et al., 1994; Davis et al., 1997; LeMaster et al., 1999; Maklad et al., 2009; O’Brien et al., 2024) and modulate neuronal activity (Mikesell et al., 2022; Moehring et al., 2018; Pang et al., 2015; Sadler et al., 2020). Our findings extend the roles of keratinocytes in controlling innervation by demonstrating that the mechanical state of the epidermis regulates nerve fiber severing and pruning.

We initially observed a striking loss of epidermal innervation following Notch activation in epidermal cells. Although Notch is a well-established regulator of epidermal differentiation (Blanpain et al., 2006; Moriyama et al., 2008), our findings suggest that altered contractility, rather than differentiation itself, is the primary driver of this phenotype. Notch activation in adult epidermis produced a rapid reduction in innervation that preceded detectable changes in differentiation-marker expression and occurred too quickly to be explained by epidermal turnover. Moreover, pharmacological inhibition of contractility largely rescued epidermal innervation. Although these experiments do not exclude a contribution from Notch-dependent changes in differentiation, they establish increased contractility as an important mediator of the denervation phenotype. The elevated contractility of NICD-expressing epidermis was unexpected because it is not a recognized feature of the spinous-cell fate promoted by Notch (Blanpain et al., 2006; Moriyama et al., 2008). This response may therefore reflect supraphysiological or prolonged pathway activation rather than the normal consequences of endogenous Notch signaling. Accordingly, phenotypes in NICD-expressing epidermis should not necessarily be interpreted simply as extensions of the physiological role of Notch in differentiation.

Our results demonstrate that the architecture of axon terminals can be sculpted by forces generated in surrounding non-neuronal cells. These findings broaden the emerging role of cell and tissue mechanics in regulating neuronal morphology. Neuronal growth and branching are influenced by substrate stiffness and mechanical tension (Pillai and Franze, 2024). During *Drosophila* metamorphosis, developmentally destabilized sensory dendrites are severed by forces associated with abdominal contractions and tissue movement (Krämer et al., 2023). In that setting, large-scale morphogenetic forces act on neurites already committed to pruning. Our findings suggest a distinct mechanism in which spatially restricted contractility within cells immediately surrounding processes can control local homeostatic severing. Whether epidermal contractility acts by directly applying mechanical stress to neuronal processes or instead activates a mechanosensitive signaling program within neurons remains unresolved. Distinguishing between these possibilities will be important for understanding whether force is the direct executor of severing or an upstream signal that initiates a neuron-intrinsic pruning program. It will also be important to understand whether contractility-induced pruning of neurons occurs in any pathologic states where tissue mechanics are known to be altered, including cancers and fibrosis.

Despite the substantial loss of epidermal innervation following increased epidermal contractility, we did not detect acute defects in touch or temperature sensation. This apparent functional resilience could reflect redundancy among sensory afferents, amplification within peripheral or central circuits, or other compensatory changes. Such robustness may be particularly important in the skin, where sensory endings are repeatedly exposed to mechanical stress and injury. Alternatively, the behavioral assays used here may not be sensitive enough to detect changes. Thus, preserved responses in these assays should not be interpreted as evidence that sensory function is entirely unaffected.

In normal epidermis, increased actomyosin contractility is concentrated in granular keratinocytes (Miroshnikova et al., 2018; Rübsam et al., 2017) at the level at which tight junctions form and sensory nerves normally terminate. Previous work implicated tight junctions in establishing this termination boundary (Takahashi et al., 2019). Our results suggest that local contractility is a separable and sufficient component of this process because increasing epidermal contractility induces nerve severing even in the absence of tight junctions. Conversely, loss of epidermal non-muscle myosin II produces pronounced abnormalities in neurite organization, most notably the extension of nerve fibers into more superficial epidermal layers. These mice also exhibit increased touch sensitivity, suggesting that the depth and pattern of epidermal innervation contribute to sensory gain. However, this hypersensitivity cannot yet be attributed solely to superficial neurite extension. Other changes in axonal branching, terminal specialization, keratinocyte mechanics, or keratinocyte-to-neuron signaling could also alter sensory responses. Together, these data support the idea of a spatial boundary for innervation that is mechanically defined.

## Materials and Methods

### Mice

All animal work was approved by Duke University’s Institutional Animal Care and Use Committee. Both male and female mice were used in this study and housed in a facility with 12-h light/dark cycles. Genotyping for all transgenic animals was performed by standard PCR on ear or tail clips. Mouse lines used in this study include K14-Cre (JAX, 018964), K5-CreER (JAX, 029155), K10-rtTA (Muroyama and Lechler, 2017), K14-rtTA(Nguyen et al., 2006), TRE-Arhgef11^CA^ (Ning et al., 2021), Rosa-NICD (JAX, 008159), RBP-J (Han et al., 2002), Myh9 flox (Jacobelli et al., 2010), Myh10 flox (Ma et al., 2009), and CD1 (Charles River, 022).

### Antibodies

For standard immunofluorescence staining, the following primary antibodies were used: rat anti-HA (Roche, Cat#11867423001, 1:500), rat anti-β4 integrin (BD Biosciences, Cat#553745, 1:500), chicken anti-GFP (Abcam, Cat#ab13970, 1:2000), rat anti-α18 (gift from Akira Nagafuchi (Yonemura et al., 2010), Kumamoto University, 1:200), rabbit anti-muscle myosin heavy chain II-A (Biolegend, Cat#909802, 1:500), rabbit anti-muscle myosin heavy chain II-B (Biolegend, Cat#909901, 1:500), rabbit anti-myosin IIC (D4A7) (Cell Signaling, Cat#8189, 1:500), rabbit anti-YAP/TAZ (D24E4) (Cell Signaling, Cat#8418, 1:200), guinea pig anti-keratin 10 (Progen, Cat#GP-K10, 1:500), rabbit anti-loricrin (gift from Colin Jamera (Jamora et al., 2004), 1:500), rabbit anti-ZO1 (Invitrogen, Cat#61-7300, 1:500), and rabbit anti-UCH-L1/PGP9.5 (Proteintech, Cat#14730-1-AP, 1:5000). F-actin was stained with phalloidin Alexa Fluor 488 (Invitrogen, Cat#A12379, 1:5000), phalloidin TRITC (Invitrogen, Cat#R415, 1:5000) and phalloidin Alexa Fluor 647 (Invitrogen, Cat#A22287, 1:400). Nuclei were labeled with Hoechst (Invitrogen, Cat#H21492, 1:2000). All Alexa Fluor conjugated secondary antibodies were purchased from Jackson ImmunoResearch and used with dilution from 1:200-1:500.

### Reagents

Tamoxifen (Cat#T5648) and doxycycline hyclate (Cat#D9891) were purchased from Sigma-Aldrich. ROCK inhibitor Y27632 (Cat#HB2297) was purchased from hello bio. Doxycycline (200 mg/kg) mouse chow (Cat#S3888) was obtained from Bio-Serv.

### Doxycycline, tamoxifen, and drug administration

Embryonic stage 0.5 (E0.5) were determined as the day a copulatory plug was observed. Tamoxifen (Sigma-Aldrich, Cat#T5648) was prepared in corn oil and administered to mice by either oral gavage or intraperitonially injection at 75 mg/kg. The treatment conditions for each experiment are indicated in the figure legends. For inducing gene expression at embryonic stages, pregnant female mice were intraperitonially injected with 24 mg/kg of doxycycline and fed on doxycycline mouse chow (Bio-Serv, Cat#S3888) on the specified embryonic stage. For inducing gene

expression in adults, mice were intraperitonially injected with 100 mg/kg of doxycycline and fed on doxycycline chow before tissue collection. The treatment duration for each experiment is indicated in the figure legends. For the rescue experiments in embryonic epidermis, pregnant dams were oral gavaged with 75 mg/kg tamoxifen (from E15.5-E17.5) and intraperitonially injected with 10 mg/kg of Y27632 (from E16.5-E18.5) before collection. For contractility inhibition experiments, adult mice were intraperitonially injected with 100 mg/kg of doxycycline and 10 mg/kg of Y27632, and then mice were sacrificed 12 hours after the treatment.

### RNA-seq sample preparation and analysis

Back skins from E17.5 embryos were dissected and treated with 1.4 U/ml dispase II (Roche, Cat# 4942078001) for 1h, at 37°C. Epidermis was separated from the dermis, and RNA was extracted using a QIAGEN RNAeasy Mini kit (QIAGEN, Cat#74104) following the manufacturer’s protocols, with DNA digested by RNase-Free DNase (QIAGEN, Cat#79254). Biological replicates were generated from independent RNA samples, isolated from three E17.5 embryos from the same dam. Samples were collected and sent for sequencing and analysis by Novogene. Gene Ontology (GO) term analysis for differentially expressed genes was performed using the GO Enrichment Analysis(Ashburner et al., 2000; The Gene Ontology et al., 2023; Thomas et al., 2022). PCA plot was generated using Clustvis (Metsalu and Vilo, 2015). Raw data is in submission to GEO. Processed data is included as Supplemental Table 1.

### Immunohistochemistry

For skin tissue preparation, back hairy skin and paw glabrous skin were dissected, fixed in 4% PFA overnight at 4°C, washed with PBS for 30 minutes at room temperature, cryoprotected in 30% sucrose overnight at 4°C, and embedded in O.C.T. (Sakura, Cat#4583). For adult back skin, hair was removed by shaving before collection. Cryosections were prepared at the following thickness: 30 µm for neuronal staining in skin tissues and 10 µm for regular skin staining. After removing OCT by washing with PBS for 10 minutes at room temperature, tissue sections were incubated in blocking buffer (3% BSA, 5% normal goat serum, 5% normal donkey serum, and 0.2% Triton in PBS) for 30 -60 minutes at room temperature. Tissues were then incubated in primary antibodies diluted in blocking buffer overnight at 4°C. Following 3 washes with PBST (0.2% Triton in PBS), samples were incubated with secondary antibodies prepared in blocking buffer for 1 hour at room temperature. Washes with PBST were repeated as before, rinsed with PBS, and mounted in anti-fade buffer (90% glycerol in PBS with 2.5 mg/ml p-Phenylenediamine (Thermo Fisher, Cat#417481000). For staining α18 and ZO1, skin tissues were directly fresh-frozen in O.C.T. after dissection (Sakura, Cat#4583). Cryosections were either fixed in 4% PFA for 10 minutes at room temperature (for α18 staining) or methanol for 2 minutes at -20°C (for ZO1 staining), blocked for 15 minutes at room temperature, incubated with primary antibodies (15 minutes for α18; 1 hour for ZO1), washed with PBST, incubated with secondary antibodies for 15 minutes, washed and mounted for imaging.

### Protein lysates preparation from mouse epidermis for western blotting

Adult paw glabrous skin was dissected and floated on 1.4 U/ml dispase II (Roche, Cat#4942078001) in PBS at 4°C for 1 hour before separating the epidermis from the dermis. The epidermal tissue was then chopped and lysed in ice-cold RIPA buffer, supplemented with 10 mM DTT, complete protease inhibitor cocktail (Roche) and phosphate inhibitor PhosSTOP (Roche). Proteins were extracted by boiling in sample buffer and analyzed by western blotting. Primary antibodies used include: rabbit anti-muscle myosin heavy chain II-A (Biolegend, Cat#909802, 1:1000), rabbit anti-muscle myosin heavy chain II-B (Biolegend, Cat#909901, 1:1000), mouse-anti-β-tubulin (Sigma, Cat#T4026, 1:1000). Secondary antibodies used include: IRDYE 680 RD donkey anti-rabbit (LICOR, Cat#926-68073, 1: 5000), IRDYE 680 RD donkey anti-mouse (LICOR, Cat#925-68070,1: 5000), Goat anti-rabbit HRP (Jackson ImmunoResearch, Cat#111-035-003, 1:1000), and Goat anti-mouse HRP (Jackson ImmunoResearch, Cat#115-035-146, 1:20000).

### Behavioral tests for skin sensitivity

Both male and female adult mice were used in assessing mechanical and thermal sensitivities. Animals were habituated in testing chambers for at least two days before conducting behavioral tests. The room temperature and humidity remained stable throughout the tests. All behavioral assessments were performed in a blinded manner. However, after inducing epidermal contractility, mutant mice exhibited obvious skin phenotypes, preventing blinding.

#### von Frey test

Mice were habituated in boxes on an elevated metal mesh floor for at least 30 minutes before testing. Mechanical sensitivity was assessed by the up-down method, using a set of von Frey filaments (Stoelting) with logarithmically increasing stiffness from 0.02 to 2.56 g. The von Frey filaments were applied perpendicularly to the plantar surface of right hind paws with a maximum application time of 6 seconds per filament. A quick paw withdrawal or licking in response to the stimulus was recorded as a positive response. Paw withdrawal threshold (PWT) was calculated using the up-down method (Chaplan et al., 1994; Xu et al., 2025). Paw withdrawal frequency (PWF) was measured with a 0.16 g von Frey filament and defined as the number of reflexive withdrawals or licking behaviors observed in 10 stimulations. The interval between two applications was at least 5 minutes.

#### Acetone evaporation test

To assess cold sensitivity, 20 µL of acetone was applied to the plantar region of right hind paws. Behavioral responses to acetone were timed immediately after the application and observed for 60 seconds(Chen et al., 2016). Responses to acetone were graded according to the following 4-point scale (Flatters and Bennett, 2004): 0, no response; 1, quick withdrawal or flick the paw; 2, prolonged withdrawal or flicking; 3, repeated flicking and licking.

#### Hargreaves test

To measure heat sensitivity (Chaplan et al., 1994; Xu et al., 2025), paw withdrawal latency was measured using the Hargreaves radiant heat apparatus (IITC Life Science). A cut off by 20 seconds was set to prevent overheating-induced skin injury. The left hind paws were tested to avoid potential effects caused by von Frey and Acetone tests on the same day.

### Imaging, quantification and statistics

Tissue sections (8-12 µm) were imaged on a Zeiss AxioImager Z1 microscope equipped with an Apotome.2 attachment, using Plan-NEOFLUAR 40X/1.3 oil objective, Plan-NEOFLUAR 63X/1.4 oil objective, and an Axiocam 506 mono camera. Tissue sections (20-30 µm) were imaged on an inverted Zeiss 780 confocal with a Plan-APOCHROMAT 20X/0.8, Plan-APOCHROMAT DIC 40X/1.4 oil objective, or Plan-APOCHROMAT DIC 63X/1.4 oil objective. Images were acquired using Zen software (Zeiss). For intensity quantification, all images within the experiment were collected with identical exposure times and settings. All image quantification analyses were performed using Fiji software.

To quantify the density of epidermal innervation, the number of nerve fibers crossing basement membrane were counted per image. Nerve fragments in the epidermis that do not cross the basement membrane were not included. The intraepidermal nerve fiber (IENF) density (Ebenezer et al., 2007; Mangus et al., 2020) was calculated and expressed as the number of nerve fibers per millimeter of the epidermal length. For dorsal hairy skin, where extensive branching complicates discrete fiber counting, epidermal innervation was quantified as the percentage of nerve surface area (PGP9.5 positive area) within the epidermis relative to the total epidermal surface area (%). Epidermal fluorescence intensity of each protein was quantified by measuring the mean fluorescence intensity within the entire epidermal region, unless otherwise indicated. Quantification of fluorescence intensity for loricrin was measured by using line scans of consistent width across the granular layers. Quantification of fluorescence intensity for cortical F-actin, myosin IIA, myosin IIB, myosin IIC, and α-18 was performed by drawing line scans across cell-cell boundaries. The maximum fluorescence intensity at each junctional peak was measured. Quantification of nuclear YAP+ cells in the epidermis was performed by measuring numbers of nuclear YAP+ cells and total cells per epidermal region. Cell aspect ratios of granular and spinous cells (in epidermal layers 3-5) were calculated by manually tracing individual cells and measuring the ratio of cell length to cell width. The shortest distance from each nerve terminal tip to the outermost granular layer in control and Myosin KO mice was measured by drawing a straight line in Fiji.

All statistical analyses were performed with GraphPad Prism 11 software and Microsoft Excel. Statistical significance was determined by performing paired *t*-tests, unpaired *t*-tests, and one-way and two-way ANOVAS with Tukey’s post hoc test. All paired samples were control and mutant animals from the same littermates. Asterisks denote statistical significance (ns: p ≥ 0.05, ∗: p < 0.05, ∗∗: p < 0.01, ∗∗∗: p < 0.001, ∗∗∗∗: p < 0.0001). Details along with sample size were indicated in figure legends. Graphical abstracts and experimental design schematics were generated using Affinity and Adobe Illustrator.

## Acknowledgements

We thank members of the Lechler Lab for comments on the manuscript, and Matt Hilton, Colin Jamora, and Akira Nagafuchi for reagents.

## Author contribution

M.L. and T.L. conceptualized the study, designed the experiments, interpreted the data, and wrote and reviewed the manuscript. M.L. performed most experiments and collected and quantified data. T.L. conducted bulk RNA-seq experiments. J.U. assisted with sample preparation and mouse management. J.X. and R.R.J. assisted in the design and interpretation of behavioral experiments, and reviewed the manuscript. T.L. acquired funding and supervised and administered the project.

## Declaration of competing interest

The authors declare no competing interests.

## Funding

This work was supported by the following grants from the NIH to TL – R01-AR083352, R01-AR035428, and R01-AR081081 – and NIH grant R01NS13182 and DoD grant W81XWH2110756 to R.-R.J.

---

**Supplementary Figure 1.**
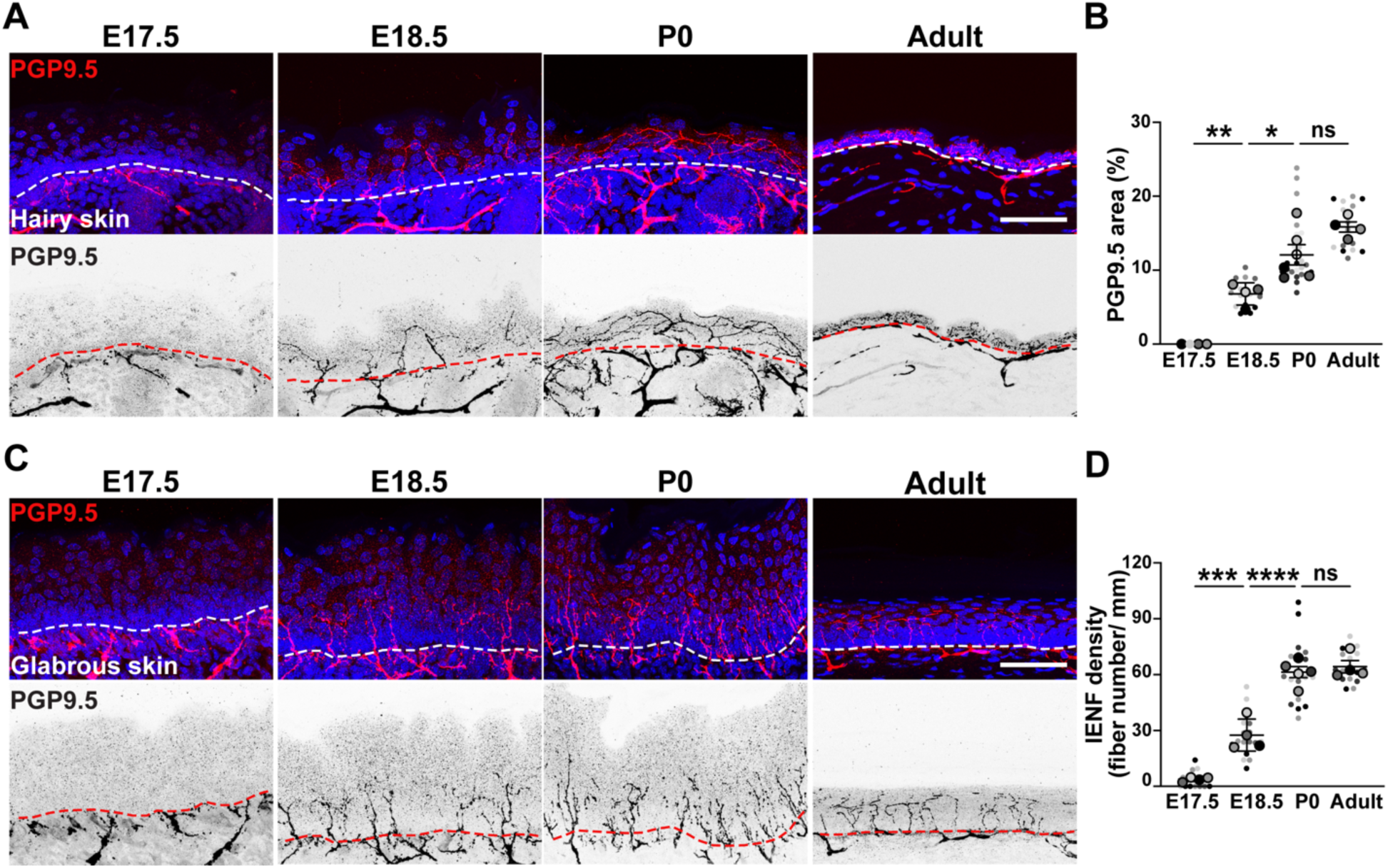
Sensory neurons innervate the epidermis after E17.5 in both back skin (hairy) and glabrous skin (non-hairy). (A) Immunofluorescence staining of nerve fibers (PGP9.5 in red; inverted image below) and nuclei (blue) in wild-type mouse back skin and (C) glabrous skin at different developmental stages. Dashed lines: basement membrane. Scale bar: 50 μm. (B) Quantification of the PGP9.5+ nerve area as a percentage of the epidermal area in wild-type mouse back skin (hairy skin). n = 4-6 animals per group, 4 fields per animal. Data are mean ± SD. ns = not significant, *p = 0.0115, **p = 0.0035, one-way ANOVA analysis. (D) Quantification of IENF density in wild-type mouse glabrous skin (non-hairy, paw skin). n = 4-5 animals per group, 3-5 fields per animal. Data are mean ± SD. ns = not significant, ***p = 0.0008, ****p < 0.0001, one-way ANOVA analysis.

**Supplementary Figure 2.**
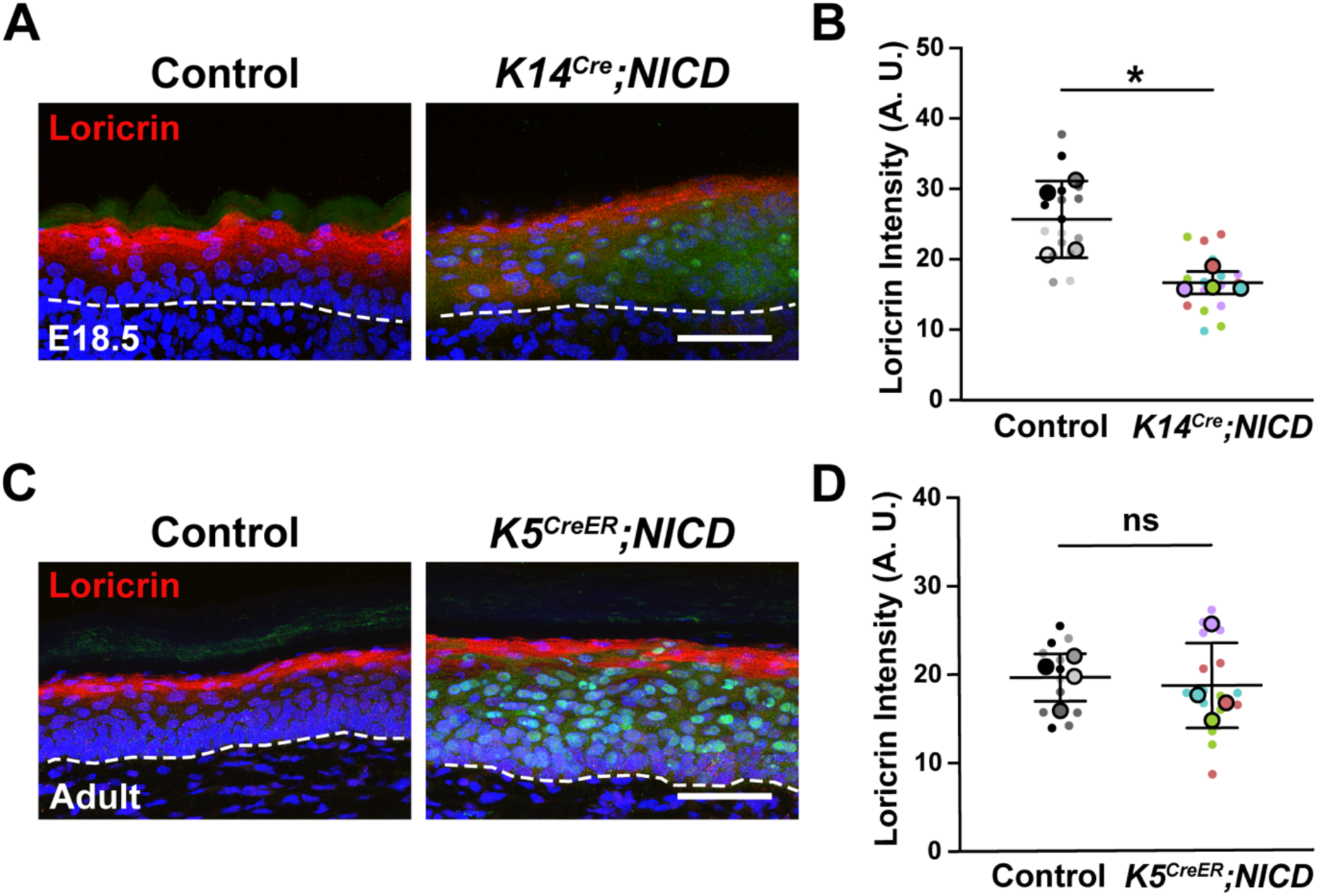
Activated Notch signaling in epidermal cells impairs epidermal differentiation during early development. (A) Immunofluorescence staining of granular marker (loricrin in red), Notch activated keratinocytes (GFP in green) and nuclei (blue) in E18.5 control and K14^Cre^;NICD back skin. Dashed lines: basement membrane. Scale bar: 50 μm. (B) Quantification of loricrin fluorescence intensity in control and K14^Cre^;NICD epidermis. n = 4 embryos per group, 4 fields per embryo. Data are mean ± SD. p = 0.337, paired t-test. (C) Immunofluorescence staining of granular marker (loricrin in red), Notch activated keratinocytes (GFP in green) and nuclei (blue) in control and K5^CreER^;NICD paw non-hairy skin in adulthood. Dashed lines: basement membrane. Scale bar: 50 μm. (D) Quantification of loricrin fluorescence intensity in control and K5^CreER^;NICD epidermis. n = 4 mice per group, 4 fields per animal. Data are mean ± SD. ns = not significant, unpaired t-test.

**Supplementary Figure 3.**
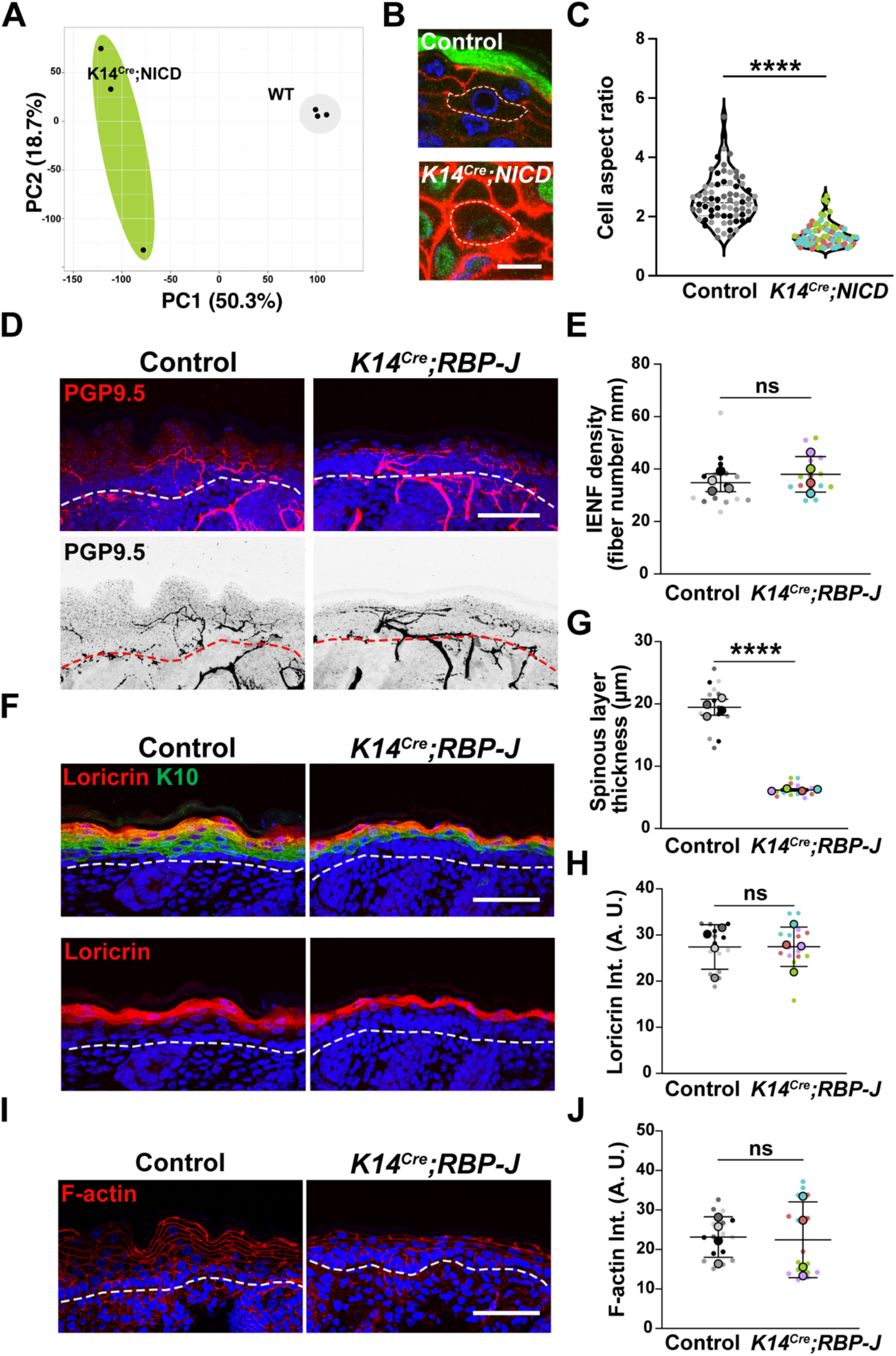
Loss of canonical Notch signaling in keratinocytes does not impair epidermal innervation. (A) A PCA plot showing the first two principal components for gene expression levels from E17.5 K14^Cre^;NICD (green) and WT samples (gray). n=3 embryos per genotype. PC1: 50.3% of the total variance. PC2: 18.8% of the total variance. (B) Magnified insets from Figure 2D. The morphology of suprabasal cells in control and K14-NICD epidermis are outlined by dashed lines. GFP in green, F-actin in red, and nuclei in blue. Scale bar: 10 μm. (C) Quantification of cell aspect ratio in control and K14-NICD epidermis at E18.5. n= 72 cells for control and n=72 cells for K14-NICD embryos. 3 embryos per genotype. Data are mean ± SD. p < 0.0001, unpaired t-test. (D) Immunofluorescence staining of neurons (PGP9.5 in red; inverted image below) and nuclei (blue) in E18.5 control and K14^Cre^;Rbp-J back skin. Dashed lines: basement membrane. Scale bar: 50 μm. (E) Diagram of the K14^Cre^;Rbp-J mouse model.(F) Quantification of IENF density in E18.5 control and K14^Cre^;Rbp-J back skin. n = 4 embryos per group, 4 fields per animal. Data are mean ± SD. ns = not significant, unpaired t-test. (G) Immunofluorescence staining of F-actin (red) and nuclei (blue) in E18.5 control and K14Cre;Rbp-J back skin. Dashed lines: basement membrane. Scale bar: 50 μm. (H) Quantification of the fluorescence intensity of F-actin in E18.5 control and K14^Cre^;Rbp-J epidermis. n = 4 embryos per group, 4 fields per embryo. Data are mean ± SD. ns = not significant, unpaired t-test. (I) Immunofluorescence staining of keratin 10 (green), loricrin (red) and nuclei (blue) in E18.5 control and K14^Cre^;Rbp-J back skin. Dashed lines: basement membrane. Scale bar: 50 μm. (J) The thickness of spinous layers (keratin 10+ and loricrin-) in control and K14^Cre^;RBP-J epidermis at E18.5. n = 4 embryos per group, 4 fields per embryo. Data are mean ± SD. p < 0.0001. (K) Quantification of loricrin fluorescence intensity in control and K14^Cre^;RBP-J epidermis at E18.5. n = 4 embryos per group, 4 fields per embryo. Data are mean ± SD. ns = not significant, unpaired t-test.

**Supplementary Figure 4.**
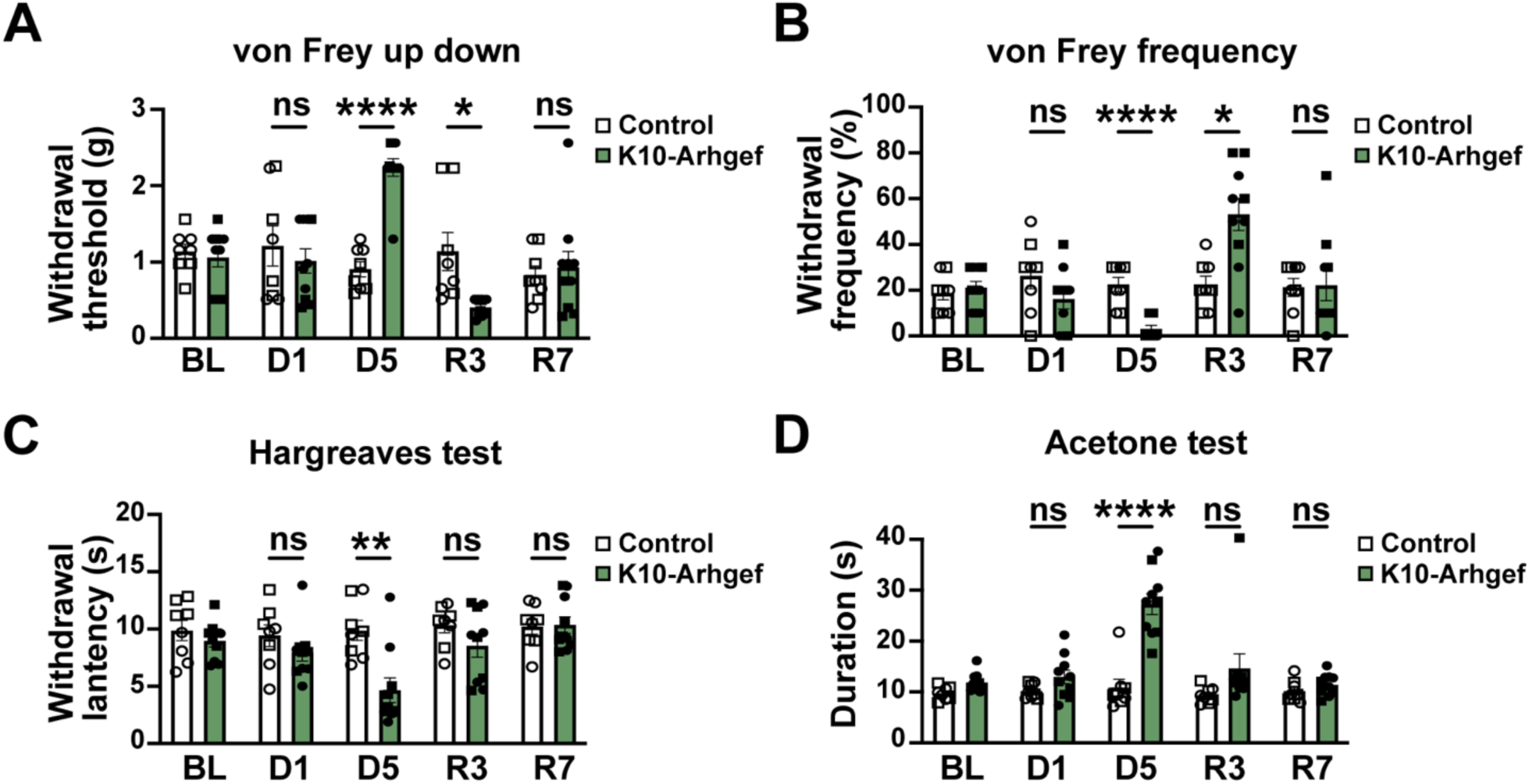
Mechanical and thermal sensitivity are restored after epidermal reinnervation. (A and B) von Frey tests showing increased mechanical sensitivity in K10-Arhgef mice after 5 days on doxycycline treatment and 3 days after doxycycline removal. n = 8-10 per genotype. Control group, 4 males and 4 females; K10-Arhgef11 group, 5 males and 5 females. Data are mean ± SEM. ns = not significant, *p= 0.0222, **p= 0.0019, ***p= 0.0002, ****p <0.0001, two-way ANOVA followed by Sidak post hoc test. Circle, male; square, female. (C and D) Hargreaves tests and acetone tests showing increased thermal and cold sensitivity in K10-Arhgef11 mice after 5 days on doxycycline treatment and returned to normal after dox removal. n = 8-10 per genotype. Control group, 4 males and 4 females; K10-Arhgef11 group, 5 males and 5 females. Data are mean ± SEM. ns = not significant, **p=0.0018, ****p <0.0001, two-way ANOVA followed by Sidak post hoc test. Circle, male; square, female.

**Supplementary Figure 5.**
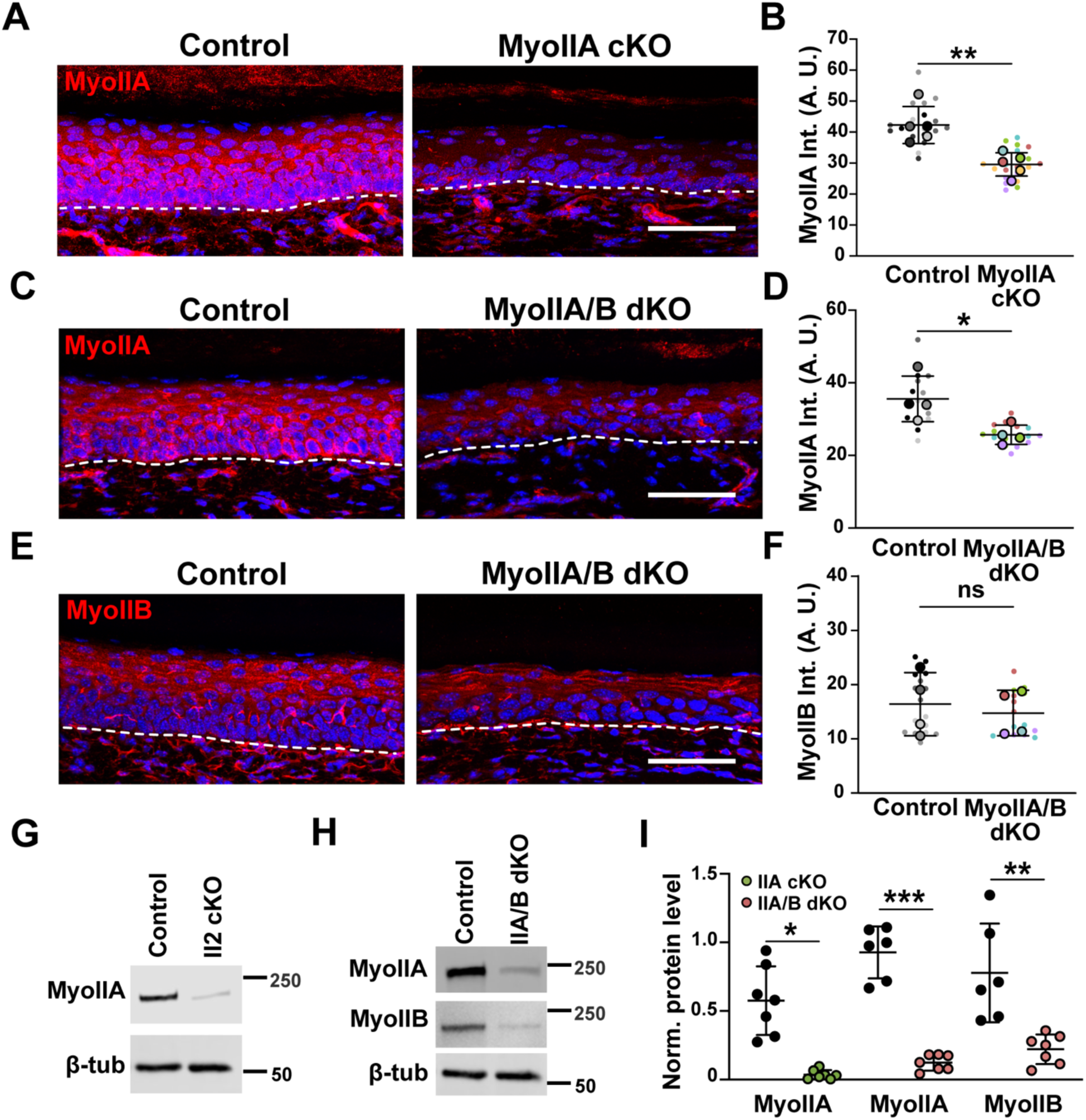
Validation of ablation of non-muscle myosin IIA and IIB proteins. (A) Immunofluorescence staining of myosin IIA in control and K5^CreER^;Myh9 adult paw non-hairy skin at 7 days post-tamoxifen. Dashed lines: basement membrane. Scale bar: 50 μm. (B) Fluorescence intensity of myosin IIA in control and MyoIIA cKO adult paw non-hairy skin. n = 5 mice per group, 4 fields per animal. Data are mean ± SD. p = 0.0038, ns = not significant, unpaired t-test. (C and E) Immunofluorescence staining of myosin IIA and myosin IIB in control and MyoIIA/B dKO adult paw non-hairy skin 7 days post-tamoxifen. Dashed lines: basement membrane. Scale bar: 50 μm. (D) Fluorescence intensity of myosin IIA and (F) myosin IIB in control and MyoIIA/B dKO adult paw non-hairy skin. n = 4 mice per group, 4 fields per animal. Data are mean ± SD. p = 0.0274. ns = not significant, unpaired t-test. (G) Western blots of myosin IIA in control and MyoIIA cKO epidermis in adult mice. (H) Western blots of myosin IIA and myosin IIB in control and MyoIIA/B dKO epidermis in adult mice. (I) Quantification of myosin IIA and myosin IIB level from (G) and (H). n = 6-7 mice per group. Data are mean ± SD. ****p <0.0001, ***p =0.001, **p = 0.0024, unpaired t-test.

**Supplementary Figure 6.**
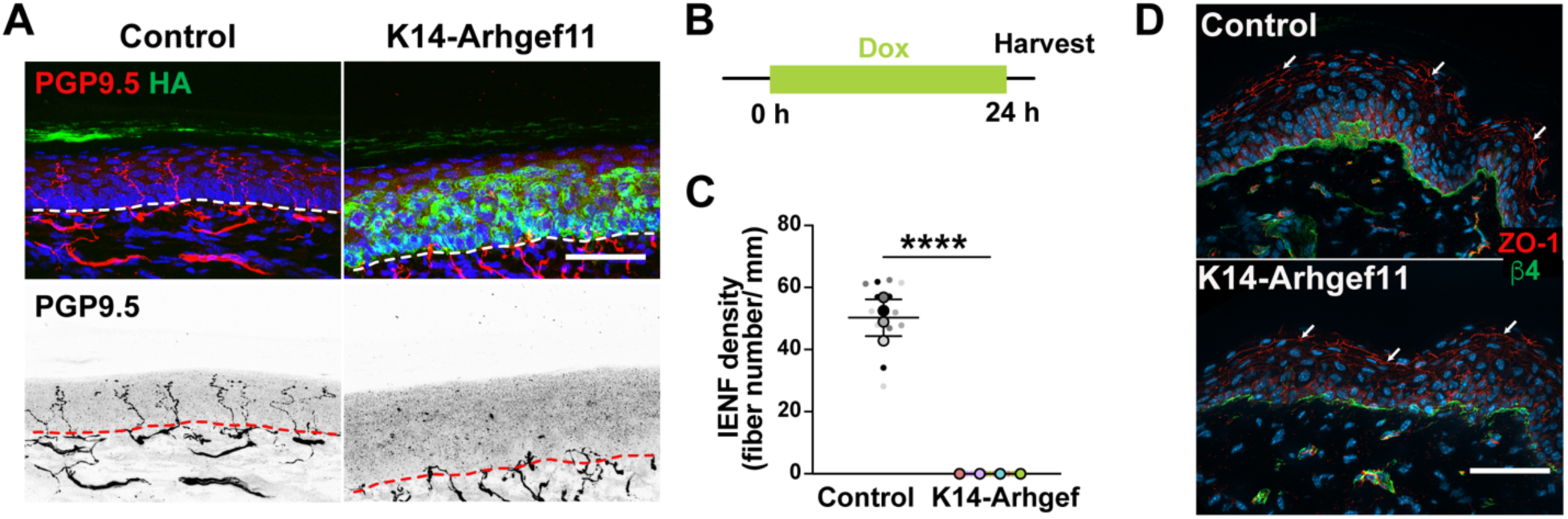
Increasing epidermal contractility does not induce ectopic tight junction assembly. (A) Immunofluorescence staining of nerve fibers (PGP9.5 in red), contractile keratinocytes (HA in green) and nuclei (blue) in control and K14-Arhgef11 adult paw non-hairy skin. Mice were treated with doxycycline for 24 hours. Dashed lines: basement membrane. Scale bar: 50 μm. Noted that the cornified envelope on the top of epidermis has autofluorescence in green. (B) Timeline of the doxycycline treatment. (C) Quantification of IENF density in adult paw non-hairy skin. n = 4 mice per group, 3-4 fields per animal. Data are mean ± SD. p < 0.0001, unpaired t-test. (D) Immunofluorescence staining of tight junction protein (ZO-1 in red), basement membrane marker (β4-Integrin in green) and nuclei (blue) in control and K14-Arhgef11 adult paw non-hairy skin. Arrows point to ZO-1 accumulation at cell borders in granular cells. Scale bar: 50 μm.

## Notes

### Competing Interest Statement

The authors have declared no competing interest.

